# Protein kinase signalling at the *Leishmania* kinetochore captured by XL-BioID

**DOI:** 10.1101/2021.07.08.451598

**Authors:** Vincent Geoghegan, Nathaniel G. Jones, Adam Dowle, Jeremy C. Mottram

## Abstract

Elucidating protein kinase signaling pathways is an important but challenging problem in cell biology. Phosphoproteomics has been used to identify many phosphorylation sites, however the spatial context of these sites within the cell is mostly unknown, making it difficult to reconstruct signalling pathways. To address this problem an *in vivo* proximity capturing workflow was developed, consisting of proximity biotinylation followed by protein cross-linking (XL-BioID). This was applied to protein kinases of the *Leishmania* kinetochore, leading to the discovery of a novel essential kinetochore protein, KKT26. XL-BioID enabled the quantification of proximal phosphosites at the kinetochore through the cell cycle, allowing the phosphorylation state of the kinetochore to be followed during assembly. A specific inhibitor of kinetochore protein kinases KKT10/KKT19 was used to show that XL-BioID provides a spatially focussed view of protein kinase inhibition, identifying 16 inhibitor-responsive proximal phosphosites, including 3 on KKT2, demonstrating the potential of this approach for discovery of *in vivo* kinase signalling pathways.

## Introduction

Phosphorylation events catalysed by protein kinases are key biochemical signals that form the basis for many important signal transduction pathways in eukaryotes, allowing the cell to coordinate complex intracellular molecular processes while integrating information about external conditions^1^. Deciphering protein kinase mediated signalling pathways is a major challenge, often requiring multiple approaches such as genetic screens, chemical inhibition and *in vitro* reconstitution^2^. Amongst a variety of approaches, phosphoproteomics has developed into a powerful method for studying phosphorylation based signalling pathways, providing quantitative information on individual phosphorylation sites. Such studies are commonly carried out on samples derived from whole lysed cells which can only provide information on total cellular levels of any particular phosphorylation site; the spatial information on where the phosphorylation site occurred within the cell is often lost^3^. Sub-cellular fractionation can be used to provide some spatial information for phosphosites, albeit at limited resolution^4^. Affinity purification of a complex of interest can also lead to identification of any phosphosites present, however these approaches rely on the spatial information surviving lysis and purification; interactions within signalling pathways are often transient and weak, representing a particular challenge.

Proximity labelling is emerging as an additional technology enabling the study of molecular complexes within the cell^5^. In biotin ligase-based proximity labelling, a protein of interest is genetically fused to a promiscuous biotin ligase. This fusion protein is expressed within the cell and upon addition of biotin the biotin ligase generates a ‘cloud’ of reactive biotin ester, proximal proteins then become covalently tagged with biotin. High resolution spatial information (10-20 nm) is thus encoded by *in vivo* biotinylation, proximal proteins are then enriched using streptavidin and identified by mass spectrometry based proteomics^6^. Since spatial information is encoded by *in vivo* biotinylation, there is no requirement to maintain complexes post-lysis. This has the significant benefit of potentially capturing more transient interactions and permitting the use of very harsh lysis conditions to extract components of structures that are difficult to solubilise. Moreover, the streptavidin-biotin interaction formed during enrichment of proximal material is very strong, allowing the use of stringent wash conditions, reducing the occurrence of post-lysis artefactual interactions.

The unique beneficial aspects of proximity biotinylation mean that it has great potential in the elucidation of protein kinase signaling pathways and has already been used to investigate mammalian mitogen activated protein kinase (MAPK), Hippo and EGFR signaling pathways^7–9^. However, such studies have not provided quantitative phosphosite level detail, likely due to the challenge of identifying and quantifying phosphorylation sites from proximal material which is limited in amount. To overcome this challenge, we have combined proximity biotinylation with *in vivo* protein cross-linking (XL-BioID), which substantially increases the amount of proximal material recovered and allows identification of proximal phosphosites.

We have applied XL-BioID to gain insight into the molecular architecture and protein kinase signalling at the kinetochore in *Leishmania mexicana,* a parasite that causes the human disease leishmaniasis^10^. The kinetochore is a large protein complex present in all eukaryotes that links chromosomes to force-generating spindle microtubules, ensuring accurate chromosome segregation during cell division. The inner kinetochore is bound to chromatin throughout the cell cycle, whilst the outer kinetochore assembles during mitosis. Intriguingly, the 25 known inner kinetochore proteins in *Trypanosoma* and *Leishmania* do not bear any readily detectable sequence homology to kinetochore components in other eukaryotes^11,12^. Another divergent feature is that four protein kinases, KKT2, KKT3, KKT10 (CLK1) and KKT19 (CLK2), are present in the core structure of the kinetochore. Studying the molecular detail of the *Leishmania* kinetochore may therefore reveal unique solutions that have evolved to solve the fundamental problem of chromosome segregation and provide information to drug discovery programs targeting this essential protein complex^13^.

We first show that XL-BioID leads to a ~10 fold increase in signal intensity for purified, proximal kinetochore proteins compared to conventional BioID, taking advantage of this to understand the molecular environment of KKT2, KKT3 and KKT19. In doing so, we discovered two novel essential components of the kinetochore, KKT24 and KKT26. To map the kinetochore as it assembles during the cell cycle we synchronised parasites expressing endogenously miniTurboID tagged KKT3 and used XL-BioID to take a spatial ‘snapshot’ at G1/S, S and G2/M phase. We show that XL-BioID can identify and quantify both proximal proteins and phosphosites from the same sample, enabling us to follow the dynamics of proximal proteins and proximal phosphosites at the kinetochore through the cell cycle, revealing proximal phosphosites with dynamic profiles distinct from that of the parent protein at the complex. Finally, we treated parasites with the specific KKT10/KKT19 inhibitor AB1, and used XL-BioID to detect a set of kinetochore proximal phosphosites that were significantly reduced in G2/M stage parasites. Quantitative information on proximal levels of proteins allowed us to distinguish between proximal phosphosites that reduced due to complex dissociation and those that reduced whilst the corresponding protein remained proximal. Amongst the latter, were 3 phosphosites on KKT2 which is a known substrate for KKT10/KKT19 in *T. brucei^14^*.

## Results

### XL-BioID increases *in vivo* capture of proximal proteins with DSP cross-linking

We sought to improve the coverage of proximity biotinylation by adding a second *in vivo* proximity capturing step. We used dithiobis(succinimidyl propionate) (DSP), a membrane permeable amine reactive cross-linker with a 12 Å spacer to covalently link proximal proteins that escaped biotinylation, we therefore call this approach XL-BioID (Fig. 1a). The covalent links withstand the harsh solubilisation and washing that are a key benefit of the proximity biotinylation workflow. Because capturing proximity information is not solely reliant on protein-crosslinking, lower concentrations of cross-linker can be used to minimise co-purification of non-proximal proteins.

**Fig. 1:**
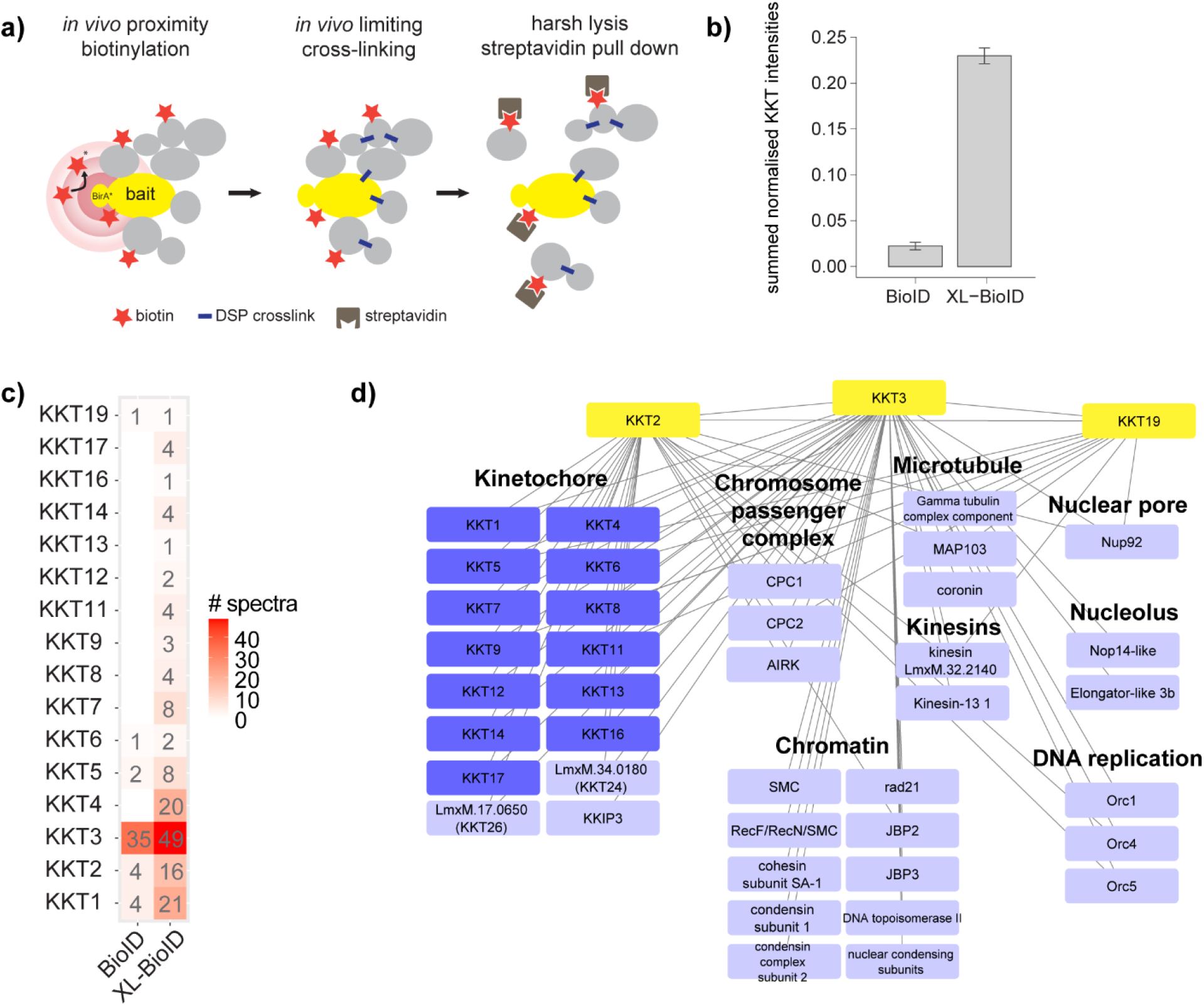
XL-BioID increases enrichment of proximal proteins. **a**, Schematic of XL-BioID. Addition of exogenous biotin initiates *in vivo* biotinylation of proteins proximal to the BirA* tagged bait protein. A membrane permeable cross-linker, DSP, is then used in a second *in vivo* proximity capturing step to covalently link any proximal proteins that escaped biotinylation. Biotin tagging and covalent linking survive harsh lysis enabling the complex to be solubilised for affinity purification. **b**, Comparison of kinetochore protein amounts purified in conventional BioID and XL-BioID. Label free intensities of kinetochore proteins (KKTs) were summed and normalised to amount of purified bait (BirA*::KKT3). Median and standard deviation are shown of 3 independent replicates. **c**, Fragmentation spectra matching KKTs in BioID and XL-BioID. Median number of spectra matching to KKTs from 3 independent replicates. **d**, KKT2, KKT3, KKT19 proximal proteins identified with XL-BioID. 98 proteins were significantly enriched compared to a control line expressing BirA*::BDF5 and are classed as proximal at 1% false discovery rate (FDR). Only those proteins with annotated functions are shown.

To demonstrate the benefits of XL-BioID, we endogenously tagged the *Leishmania* kinetochore protein kinase KKT3 with the promiscuous biotin ligase BirA* using Cas9^15^. We then performed proximity biotinylation with and without the second cross-linking step, assessing the number and label free intensity of kinetochore proteins identified by mass spectrometry. The sum of kinetochore protein label free intensities were normalised to the bait intensity, resulting in a normalised label free intensity of 0.23 which was ~10 fold higher than in conventional BioID (0.022, Fig. 1b). The increased amount of proximal material allowed the identification of 16 out of the 25 inner kinetochore proteins in XL-BioID against 6 from BioID (Fig. 1c, Supplementary Table 1).

### Proximal environment of kinetochore kinases captured by XL-BioID

A distinct feature of kinetoplastid kinetochores is the presence of protein kinases apparently as core components^11^. To understand the proximal environment of these kinases, we used BirA* tagged KKT2, KKT3 and KKT19 and XL-BioID to identify proximal proteins. Identified proteins were quantified by precursor intensity-based label free quantification to calculate enrichment against a control sample derived from parasites expressing a BirA* tagged bromodomain containing protein BDF5, which also localises to the nucleus, but is primarily found on chromatin at sites of transcription initiation/termination and thus serves as a spatial reference. From the 3 baits, a total of 98 high confidence proximal proteins were identified at 1% FDR (Fig. 1d and Supplementary Tables 2 and 3). KKT2 and KKT3 were associated with 59 and 63 proximal proteins respectively, compared to 26 for KKT19, likely due to the constitutive localisation of KKT2 and KKT3 to the kinetochore, whilst KKT19 appears to localise to the kinetochore more transiently during mitosis^11,16^. This is also reflected in the number of kinetochore proteins captured by each bait, KKT2 and KKT3 co-enrich 15 other kinetochore proteins, but KKT19 only 5. Gene ontology enrichment analysis of proximal proteins identified ‘kinetochore’ and ‘chromosome, centromeric region’ as the most significantly enriched terms, indicating the BirA* tagged kinases were sampling the intended set of proximal proteins. Chromosome segregation is driven by the mitotic spindle and spindle associated proteins such as kinesins, over-representation of the GO terms ‘spindle’ and ‘spindle pole’ indicate these structures became labelled as chromosomes segregated during mitosis. An important group of spindle associated proteins, the chromosome passenger complex, was identified, including the *Leishmania* orthologue of human Aurora-B kinase, AIRK, which is required for assembly of the mitotic spindle in trypanosomes ^17 18.^

KKT2 and KKT3 are constitutively bound to chromatin at the centromere, which is the assembly site for the kinetochore^11,19^. We detected a number of proximal proteins with activities in regulation of chromatin structure such as a rad21, cohesin subunit, condensin subunits 1 and 2, SMC and nuclear condensing subunits. A key molecule that epigenetically specifies a region of chromatin as the centromere is the centromeric histone CENP-A. *Leishmania* and other kinetoplastids are notable in lacking this histone variant, raising the question of how the centromere is defined in these eukaryotes. We found JBP2 proximal to KKT2 and KKT3, JBP3 was also found proximal to KKT3. JBP2 and JBP3 are involved in synthesis and binding of base-J respectively, a glycosylated thymidine DNA base found only in kinetoplastids^20^. Base-J has been shown to be enriched at centromeres^21^, the finding that JBP2 and JBP3 are close to the kinetochore raise the possibility of base-J having a role in specifying the kinetochore binding site in *Leishmania.* The centromere in *Leishmania* is also the major origin of replication for chromosomes, explaining the enrichment of the origin of replication proteins Orc1, Orc4 and Orc5^22^. After replication of DNA, cohesion of sister chromatids is established, requiring the cohesin complex, which has been shown to accumulate at centromeres in budding yeast^23^. Sister chromatids may also be held together by DNA catenation, which also appear to be concentrated at centromeres, until the resolving action of DNA topoisomerase II at metaphase^24^. We found both cohesin and DNA topoisomerase II to be proximal to the kinetochore, evidence that similar mechanisms underlie chromatid cohesion in *Leishmania*. Thirty of the 98 proximal proteins have no annotated function and may represent novel factors involved in chromosome segregation (Supplementary Table 2).

### KKT26 is a novel essential *Leishmania* kinetochore component

Our XL-BioID analysis of the *Leishmania* kinetochore protein kinases identified a group of unknown proximal proteins. To determine if any of these were previously undescribed components of the kinetoplastid kinetochore, we endogenously tagged uncharacterised proteins with mNeonGreen and performed a fluorescence co-localisation screen in parasites expressing mCherry KKT2. This revealed 2 previously unknown components of the kinetochore, LmxM.34.0180 and LmxM.17.0650. LmxM.34.0180 has since been identified as kinetochore protein KKT24 in *Trypanosoma brucei^12^* and we designate LmxM.17.0650 as novel kinetochore protein KKT26. Both kinetochore proteins localise clearly to the kinetochore from S phase onwards, appearing first as a characteristic set of multiple nuclear punctae which then appear to coalesce into mostly individual puncta at metaphase and anaphase (Fig. 2a). Interestingly, KKT24 also localises to the kinetoplast, a specialised region containing the mitochondrial DNA.

**Fig. 2:**
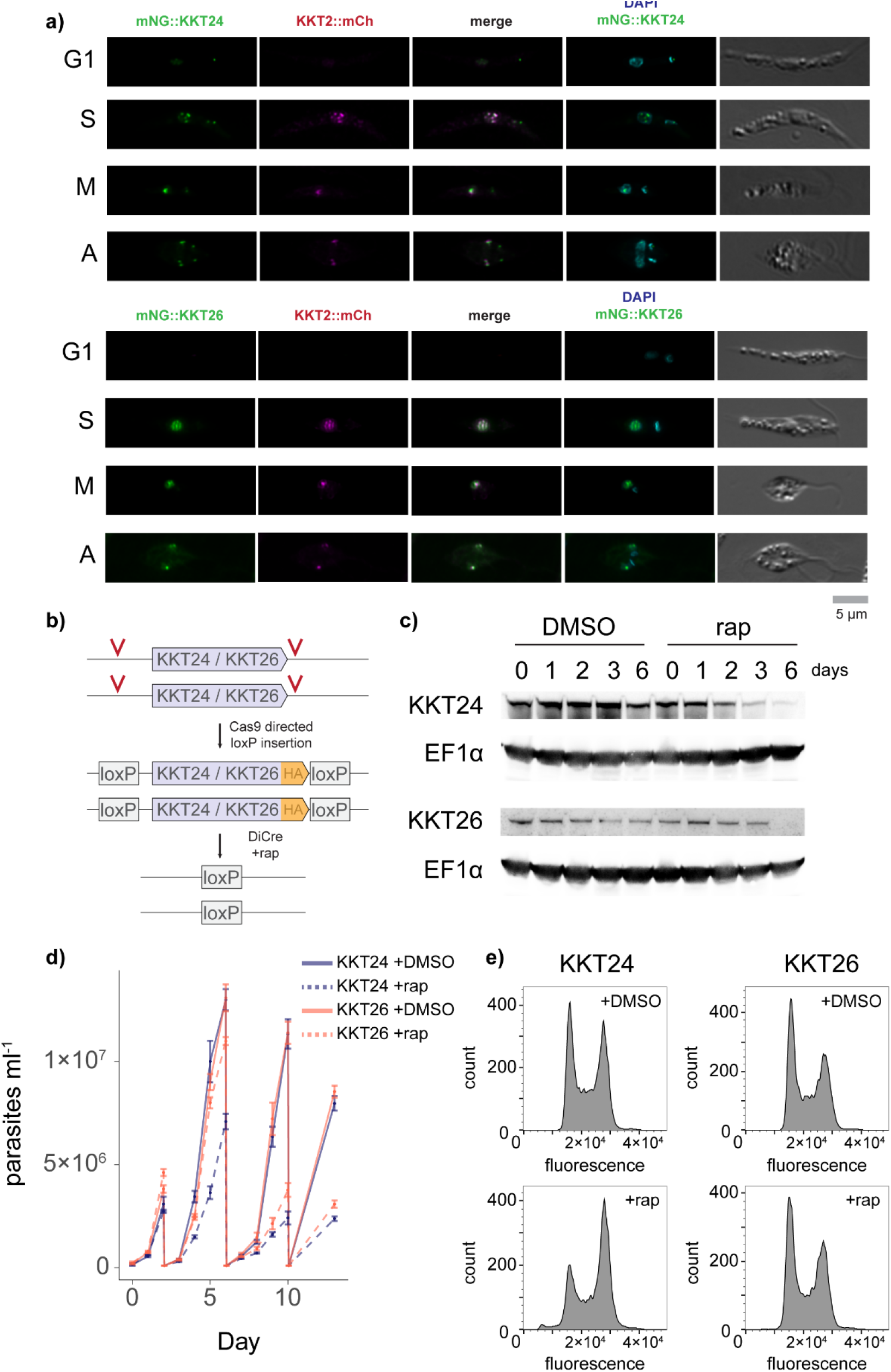
XL-BioID identifies KKT26, a novel essential component of the *Leishmania* kinetochore. **a**, Fluorescence co-localisation microscopy of KKT24 (LmxM.34.0180), KKT26 (LmxM.17.0650) with KKT2. KKT24 and KKT26 were mNeonGreen (mNG) tagged and localisations compared to mCherry (mCh) tagged KKT2 at different phases of the cell cycle: G1, S, M (metaphase), A (anaphase). **b**, Strategy for inducible deletion of KKT24 or KKT26. Homology directed repair after Cas9 induced breaks was used to insert LoxP sites flanking the endogenous genes at both alleles. Addition of rapamycin induced dimerisation of DiCre recombinase which excised KKT24 or KKT26. **c**, Depletion of KKT24 or KKT26 protein following gene excision, shown by Western blotting using anti-HA antibody. EF1ᵅ was used as a loading control. **d**, Parasite growth after KKT24 or KKT26 excision. Parasites were diluted to 1 × 10^6^ ml^−1^ at days 2, 6 and 10 after addition of rapamycin (rap) or DMSO, median and standard deviation of 4 replicates are shown. **e**, Cell cycle profile of parasites after KKT24 or KKT26 excision. Parasites 6 days after addition of rapamycin or DMSO were stained with propidium iodide for DNA content and analysed by flow cytometry.

Attempts to knockout KKT24 or KKT26 by Cas9 in *L. mexicana* promastigotes were unsuccessful, indicating these kinetochore proteins are essential to parasite survival. We therefore opted for an inducible gene deletion strategy, using Cas9-directed insertion to introduce loxP sites flanking the endogenous genes^25^. Addition of rapamycin induces dimerisation of DiCre recombinase which then excises the loxP flanked gene (Fig. 2b). We monitored protein levels after gene excision and found that KKT24 became depleted from day 3 whilst KKT26 levels depleted more slowly (Fig. 2c). We found that while both proteins are required for parasite growth, the onset of growth inhibition was earlier after KKT24 excision, matching the more rapid protein depletion kinetics compared to KKT26 (Fig. 2d). Because the kinetochore is essential for chromosome segregation and normal cell cycle progression, we measured DNA content of parasites by flow cytometry 6 days after KKT24 or KKT26 excision. Deleting KKT26 did not lead to defects in the cell cycle profile, by contrast deletion of KKT24 caused parasites to accumulate in G2/M phase and generated parasites containing sub genome equivalents of DNA (Fig. 2e). This indicates that KKT24 is important for a kinetochore activity that has an associated mitotic checkpoint.

### Proximal phosphosites are identified by XL-BioID during kinetochore assembly

Our discovery that XL-BioID purifies ~10 fold more proximal material than the standard BioID workflow suggested to us that it may be possible to enrich phosphosites from proximal proteins isolated by XL-BioID. We designed a workflow which uses XL-BioID to enrich proximal proteins that are digested on streptavidin beads to release the peptides. 60% of this is then used to enrich for proximal phosphopeptides using a Ti-IMAC resin designed for highly sensitive phosphopeptide enrichment. The remaining 40% of peptides are used as the ‘Total’ proximal sample, thus from the same sample both proximal proteins and their phosphosites can be quantified (Fig. 3a). As there is minimal knowledge on how the *Leishmania* kinetochore assembles or is regulated by phosphorylation during cell cycle progression, we used XL-BioID to follow KKT3 proximal proteins and phosphosites in synchronised parasites as they progress through the cell cycle. To achieve this, we employed the engineered BirA variant, miniTurboID, which enables proximity biotinylation at a time resolution of 10 mins compared to 18-24 hours for BirA*^26^. We were able to endogenously tag both alleles of KKT3 with miniTurboID, since KKT3 is an essential protein kinase in *Leishmania* this indicated that the tag did not block protein function^27^. Parasites were synchronised with hydroxyurea at early S phase, then released to resume the cell cycle ^21^. A 30 min biotinylation was performed at 0, 4 and 8 hrs following hydroxyurea release, which samples KKT3 proximal proteins and phosphosites at G1/S, S and G2/M stages respectively. At each timepoint, parasites were analysed by flow cytometry, confirming synchronous progression through the cell cycle (Fig. 3b). To detect proximal proteins and phosphosites with XL-BioID, label free quantitation was used and enrichment was calculated relative to samples from parasites expressing a miniTurboID tagged nuclear bromodomain protein BDF5 (Supplementary Tables 4 and 5).

**Fig. 3:**
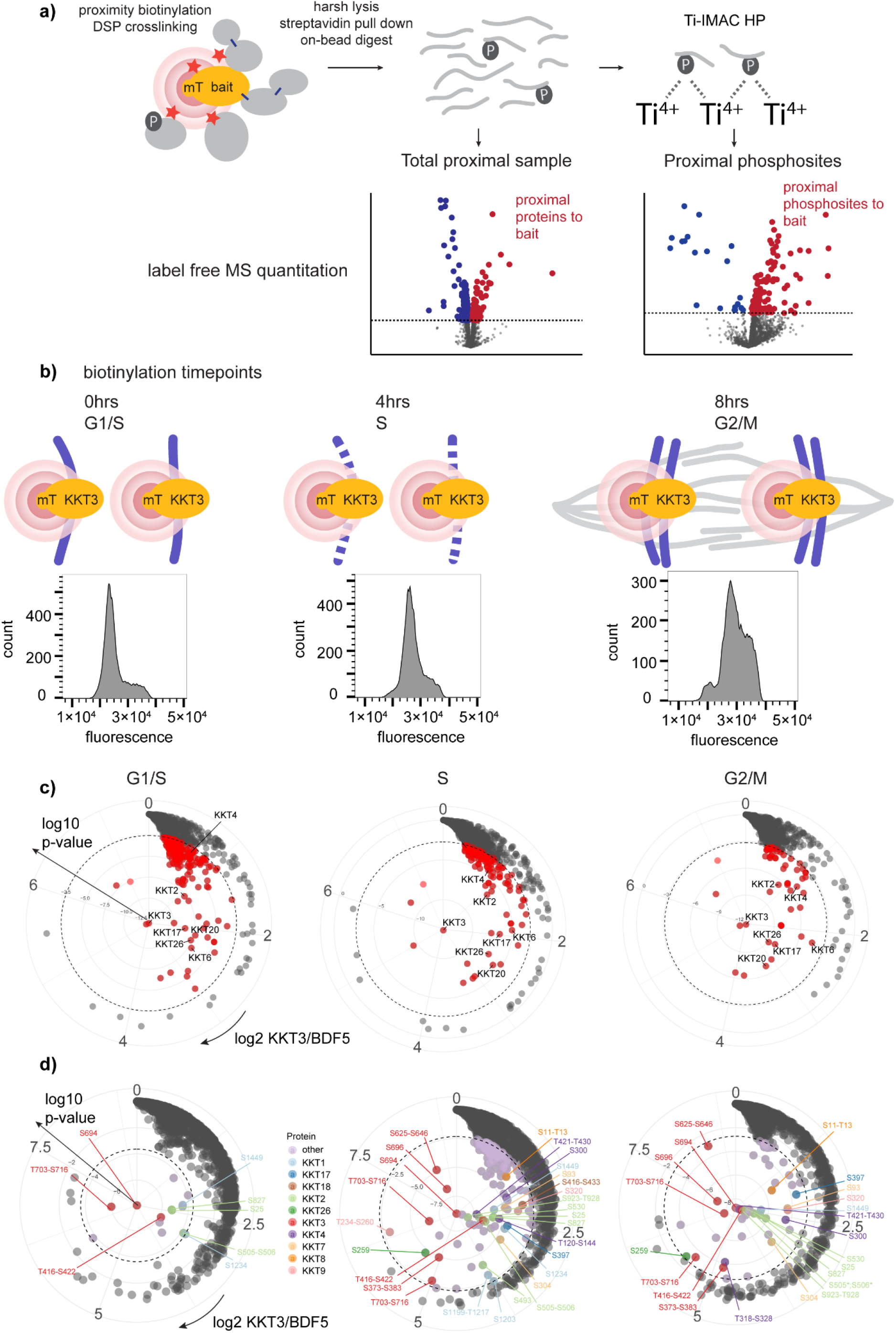
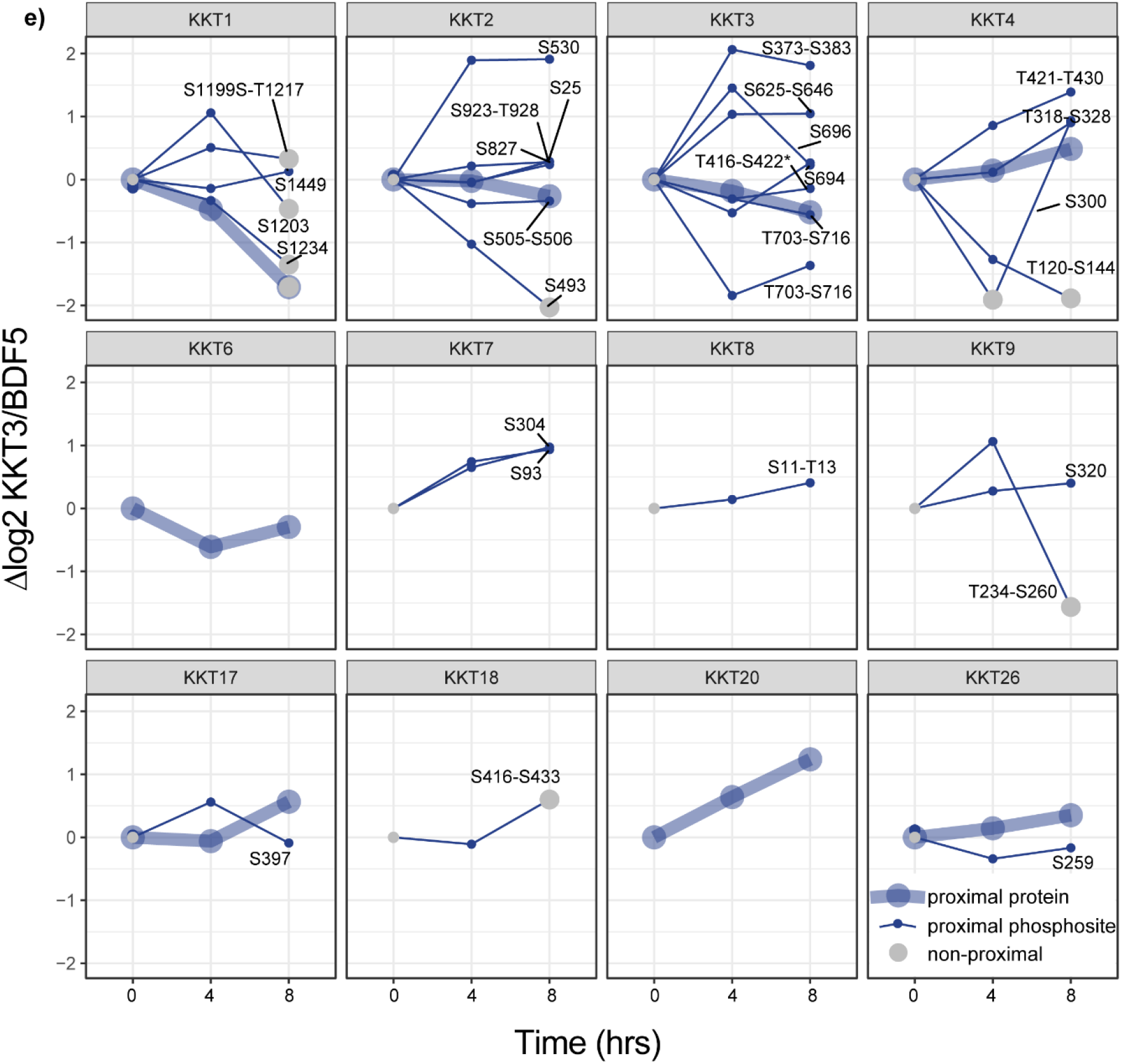
Dynamics of proximal proteins and phosphosites during kinetochore assembly revealed by XL-BioID. **a**, Schematic of XL-BioID workflow to quantify proximal proteins and proximal phosphosites from the same sample. Proximity biotinylation and DSP protein crosslinking captures proximal proteins which are then on bead digested to peptides for protein and phosphopeptide identification. Proteins or phosphosites significantly enriched (red dots) compared to a control are classed as proximal. **b**, Using KKT3::mT as a bait and XL-BioID to follow kinetochore assembly during the cell cycle in synchronised parasites. A 30 mins biotinylation ‘snapshot’ was taken at 3 timepoints after release from hydroxyurea synchronisation. Histograms of parasite DNA content are shown at each timepoint. Dotted line represents replicating chromosomes **c**, Radial plots of proteins enriched in KKT3::mT samples compared to control BDF5::mT samples at each timepoint. Log2 fold enrichment increases in a clockwise direction, log10 p-values increase from the centre outwards. Dashed line indicates 1% FDR cutoff for distinguishing proximal (red points) and non-proximal (grey points) proteins. Label free quantification was performed on 5 independent biological replicates. **d**, Radial plots of phosphosites enriched in KKT3::mT samples compared to control BDF5::mT samples at each timepoint. Dashed line indicates 5% FDR cutoff for distinguishing proximal (coloured points) and non-proximal (grey points) phosphosites. Label free quantification was performed on 5 independent biological replicates. Where a phosphosite could not be confidently localised, the possible phosphorylated region is indicated. **e**, Proximal profile plots of kinetochore proteins and phosphosites during the cell cycle. Kinetochore proteins and phosphosites proximal in at least one of the timepoints are shown. At each timepoint, mean log2 fold enrichments against BDF5::mT control are normalised to the log2 fold enrichment at time 0, *n* = 5. Where a phosphosite could not be confidently localised, the possible phosphorylated region is indicated.

The bait itself, KKT3, was the most significantly enriched protein at all time points and is therefore the central point in radial plots of quantified proteins (Fig. 3c). Because the extent of enrichment of a protein is not only dependent on proximity, but also copy number at the complex and the number of lysine residues available for biotinylation, relative distances of proteins to the bait cannot be determined from proximity labelling data. However, it is possible to infer changes at the kinetochore complex by comparing enrichment between timepoints of the cell cycle. We found that a set of kinetochore proteins consisting of KKT1, KKT2, KKT4, KKT6, KKT17, KKT20 and KKT26 were proximal to KKT3 early in the cell cycle at G1/S (Fig. 3c left plot). As the parasites progressed through the cell cycle, the enrichment of KKT2, KKT6, KKT17 and KKT26 remained relatively constant, indicating that these proteins may form part of the stable chromatin bound inner kinetochore (Fig. 3c middle and right plots). KKT2 and KKT17 have indeed been shown to be constitutively present at the kinetochore in *T. brucei*. By contrast, our proximity data showed that KKT4 and KKT20 became 1.4-fold and 2.4-fold more enriched respectively, through the cell cycle. KKT4 is the only kinetoplastid kinetochore protein demonstrated to bind microtubules and therefore likely plays an important role in attaching the kinetochore to the mitotic spindle^28^. This process takes place later in the cell cycle, which is consistent with our proximal data showing KKT4 reaching peak enrichment at G2/M (Fig. 3c right plot). KKT20 showed the largest increase in proximal enrichment through the cell cycle which is consistent with fluorescence localisation data for this protein from *T. brucei^29^*. In contrast, KKT1 was proximal at G1/S and S phase, but not at G2/M phase. Microscopy data suggest KKT1 remains localised to the kinetochore throughout the cell cycle, a discrepancy that could be due to such studies being limited in resolution to ~300 nm, whilst proximity biotinylation can report on localised re-arrangements at a much higher spatial resolution. Overall, the set of KKT3 proximal proteins decreases in complexity through the cell cycle and may reflect the fact that the kinetochores are proximal to origins of replication which have the highest activity at G1/S.

Proximal phosphosites to KKT3 show a different trend during the cell cycle, being a limited set at G1/S and peaking in diversity at S phase. The most significantly enriched phosphosite at all time points was S694 on the bait itself (Fig. 3d). At G1/S the only detected phosphosites proximal to KKT3 were KKT1 (S1449, S1234) and KKT2 (S25, S505, S827) suggesting the kinetochore was in a hypophosphorylated state (Fig. 3d, left plot). By S phase, the phosphorylation landscape surrounding KKT3 had dramatically changed, proximal phosphosites could now be identified on KKT1-KKT4, KKT7-9, KKT17-18 and KKT26 as well as a set of non-kinetochore proteins. The majority of these kinetochore phosphorylation sites remained proximal at G2/M, whilst the number of non-kinetochore proximal phosphosites had reduced.

### Dynamics of kinetochore phosphorylation during the cell cycle

Since both protein level and phosphosite level proximal information is collected from the same sample in our XL-BioID workflow, we combined both sets of quantitative data to visualise changes in subunit quantity and phosphorylation state at the kinetochore complex during cell cycle progression (Fig. 3e). This permitted us to distinguish different classes of phosphosites, those that changed in abundance in a similar manner to the protein, and more dynamic phosphosites whose levels were not simply coupled to the amount of protein at the complex. Examples of the former class are KKT1(S1234), KKT2(S25, S505, S827, S923-T928) and KKT4(T318-S328) as their abundance profile matched the protein profile. Examples of the latter class include KKT2(S493, S530) and KKT4(T120-S144, S300, T421-T430). Because these phosphosites show dynamic behaviour de-coupled from the parent protein, we hypothesise that these sites may be particularly important in the regulation of kinetochore activity, for example by controlling the recruitment of other proteins. We noted that for KKT7, KKT8, KKT9 and KKT18 we could detect proximal phosphosites but not proximal protein in the ‘Total’ sample, indicating that these proteins are only enriched enough for their proximity to be detected after phosphopeptide purification.

### XL-BioID and chemical inhibition reveals KKT10/KKT19 dependent phosphorylation at the kinetochore

In other eukaryotes, multiple protein kinase mediated signalling pathways direct the assembly of the kinetochore and regulate its attachment to spindle microtubules. Analogous pathways likely exist in *Leishmania,* however they remain to be characterised. In the related *T. brucei*, a pair of highly similar kinetochore kinases, KKT10/KKT19, are required for progression from metaphase to anaphase^14,16^. These protein kinases are unique to kinetoplastids and are attractive drug targets, motivating recent work which identified a specific covalent inhibitor of KKT10/KKT19 called AB1^13^. Recent work identified KKT2 as a KKT10 (CLK1) substrate in *T. brucei^14^*. Having successfully used XL-BioID to discover cell cycle dependent changes in phosphorylation of the kinetochore, we then used XL-BioID together with inhibition by AB1 to attempt to map KKT10/KKT19 dependent phosphorylation at the kinetochore. To do this, parasites expressing miniTurboID tagged KKT3 were synchronised with hydroxyurea, released and treated with 60nM AB1 or DMSO. A 30min biotinylation was carried out at 0hr, 4hrs and 8hrs after hydroxyurea release and proximal proteins and phosphosites were quantified as before (Fig. 4a, Supplementary Tables 4 and 5)). Principle Component Analysis (PCA) of quantified phosphosites shows two well separated groups of samples, demonstrating that the miniTurboID tagged control protein BDF5 and miniTurboID tagged KKT3 sample a different set of proximal phosphosites (Fig. 4b). Each of these groups is further divided into 3 distinct groups corresponding to the 3 timepoints in the cell cycle, AB1 treated samples cluster closely with corresponding untreated samples indicating that only a subset of identified phosphosites were affected by AB1. Clustering BDF5 or KKT3 proximal phosphosites by their intensity profiles across the samples also shows that the phosphorylation ‘landscape’ local to each protein is quite distinct, with a much larger group of phosphosites proximal to KKT3 (Fig. 4c).

**Fig. 4:**
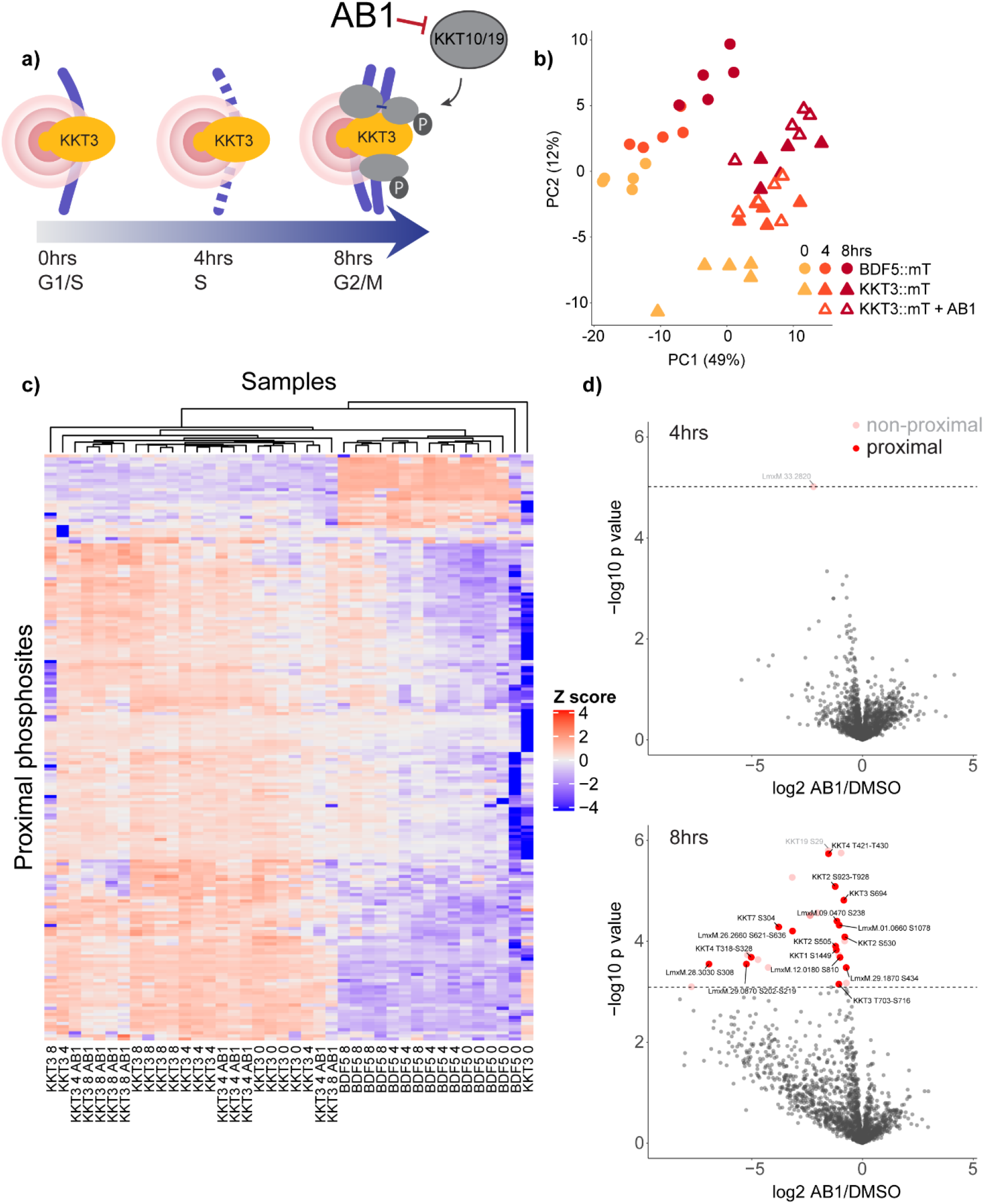
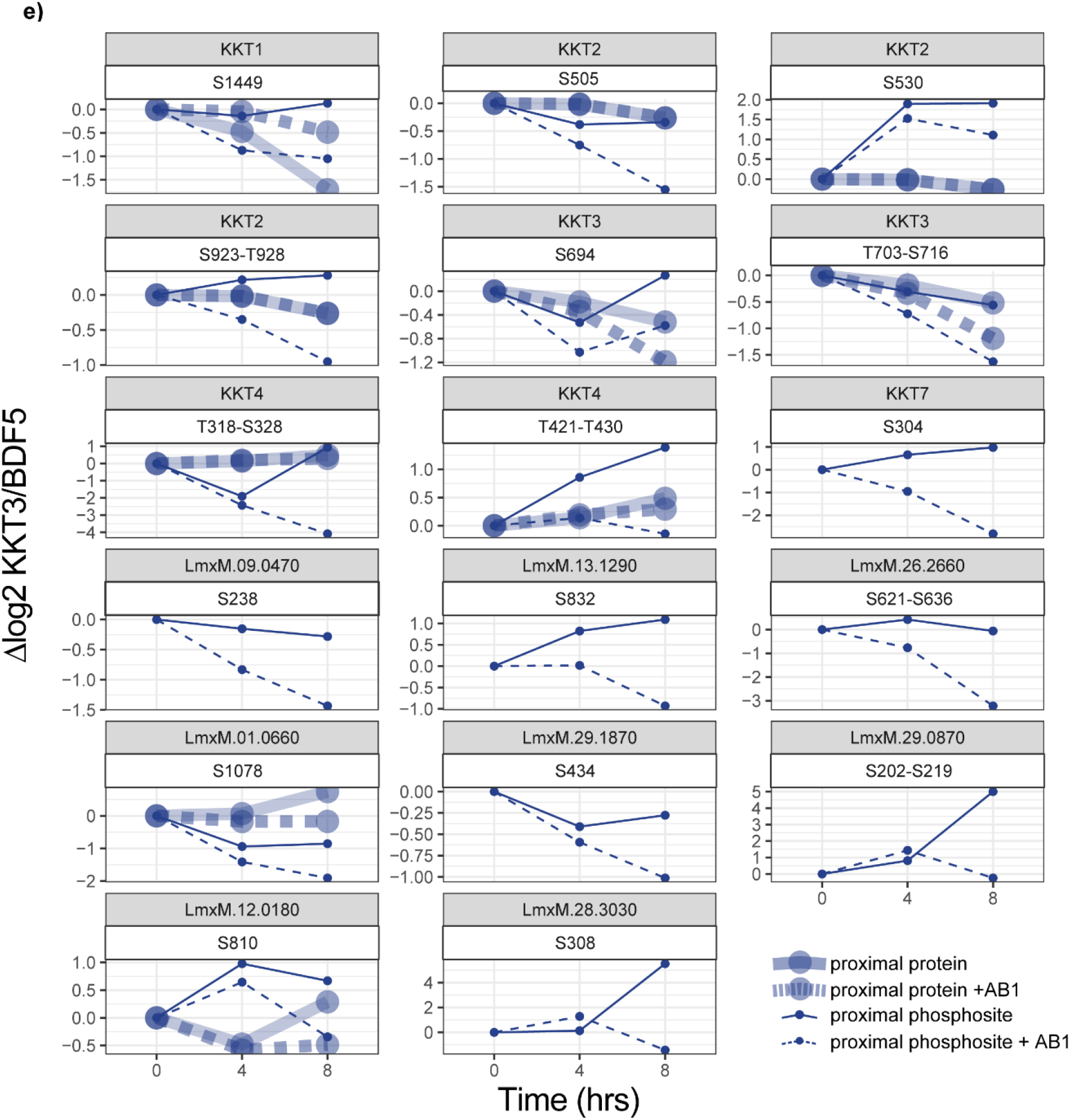
XL-BioID detects kinetochore proximal phosphosites reduced after AB1 inhibition of the kinetochore kinases KKT10/19. **a**, Experimental design as in Fig. 3 but with AB1 treatment from 0 hrs after synchronisation release. AB1 is a specific inhibitor of KKT10 and KKT19, which are required for completion of mitosis. **b**, Principal Component Analysis (PCA) of phosphopeptides quantified in XL-BioID. PCA was performed on label free quantified phosphopeptides with an ANOVA q-value < 0.1. **c**, Heatmap of proximal phosphosites. Each row is a phosphosite proximal to KKT3 or the control BDF5 in at least one timepoint at 5% FDR. Phosphosites are clustered by label free abundance similarity across the samples. Samples are also clustered by similarity. **d**, Label free quantification of all phosphopeptides identified by XL-BioID in AB1 treated vs. DMSO treated parasites at 4hrs after AB1 treatment and synchronisation release (top) and 8hrs (bottom). Significantly changing phosphosites after AB1 treatment are either non-proximal to KKT3 (pink) or proximal to KKT3 (red). Dashed line indicates 5% FDR, *n* = 5. **e**, Proximal profile plots of AB1 responsive phosphosites and protein levels. At each timepoint, mean log2 fold enrichments against BDF5::mT control are normalised to the log2 fold enrichment at time 0, *n* = 5. Where a phosphosite could not be confidently localised, the possible phosphorylated region is indicated.

To reveal the individual proximal phosphosites affected by AB1 inhibition of KKT10/KKT19, the label free intensity of phosphopeptides was compared between AB1 treated and DMSO treated parasites at 4hrs and 8hrs. At 4hrs, only one phosphosite on a regulatory subunit of protein kinase A (LmxM.33.2820) was reduced. But by 8hrs, a set of phosphosites that decrease after AB1 treatment became apparent (Fig. 4d). Because in these inhibition experiments we record an additional layer of information, the proximity to KKT3, we further screened AB1 affected phosphosites to determine those that are at or near the kinetochore. This allowed us to define 16 proximal phosphosites reduced by AB1, 9 of which were on the kinetochore itself (Fig. 4e). The most statistically significant decrease in phosphorylation occurred at S29 on KKT19, however this phosphosite was not sufficiently enriched in the KKT3 sample to be classed as proximal. Of the proximal phosphosites affected by AB1, 3 occurred on KKT2, (S505, S530, S923-T926), which is a known substrate of KKT10/KKT19 in *T. brucei^14^*, 2 occurred on KKT4 and 1 on KKT7, which have both previously been shown to be phosphorylated in a KKT10/KKT19 dependent manner in *T. brucei^16^*. For the significantly reduced proximal phosphosites, we determined if there was any change in the levels of the corresponding proximal protein. KKT2 levels at the kinetochore were unchanged by AB1 treatment, thus phosphosite decreases in this case reflect reductions in the phosphorylation state of kinetochore localised protein. Proximal KKT4 protein levels also do not change significantly after AB1 treatment, but 2 phosphosites, at T318-S328 and T421-T430, decrease significantly. Phosphorylation of KKT1 at S1449 is decreased after AB1 treatment, however proximal KKT1 levels are significantly increased, thus together indicating a large decrease in the proportion of S1449 phosphorylated KKT1 at the kinetochore. We observed that AB1 caused a decrease in levels of KKT3 by 0.63 fold at 8hrs and a similar decrease in phosphorylation sites. Since this is the bait protein, it represents the total cellular pool of KKT3. Because the amount of KKT1, KKT2 or KKT4 enriched at 8hrs is not reduced by AB1, this is evidence that the amount of KKT3 at the kinetochore is also unaffected; the decrease observed is likely to be due to free levels of KKT3 being lower in AB1 treated parasites.

## Discussion

We have developed XL-BioID, a two-step *in vivo* proximity capturing method that uses proximity biotinylation and chemical cross-linking to reveal the molecular environment around a protein of interest. Proximity labelling is being rapidly adopted as a powerful method providing high resolution spatial information between molecules. We show that by adding an orthogonal *in vivo* proximity capturing step in the form of DSP cross-linking we can recover additional proximal proteins, increasing the amount of proximal proteins enriched approximately tenfold. Although both proximity capturing steps target lysine residues, biotinylation occurs by reaction with a locally generated mixed anhydride (biotinyl-5’-AMP), whilst the subsequent cross-linking is a reaction with an exogenously added N-hydroxysuccinimide ester (DSP). The increase in proximal capture efficiency is therefore likely due to the combination of different reaction chemistries and differing local concentrations of reagent. Proximity labeling and chemical cross-linking have been previously applied to investigate the human nuclear envelope interactome, however, cross-linking was performed post-lysis after biotinylated proteins had already been purified^30^.

Importantly, we show that XL-BioID extends the capability of proximity biotinylation to quantification of phosphorylation events on proximal proteins. Phosphorylation sites have been extensively catalogued in a range of organisms by phosphoproteomics^31^. However, information on the spatial localisation of phosphorylated proteins is largely missing from such resources and is key to understanding how phosphosites are related to each other in signalling pathways. Recent work has described the identification of phosphosites in proximity biotinylation experiments using a photo-caged ER targeted biotin ligase, but this only provided organelle level spatial resolution and did not quantify proximal proteins and their phosphosites in the same sample which can limit quantification precision^32,33^. We used XL-BioID to study components of the *Leishmania* kinetochore and closely associated complexes, as well as providing new insights into protein kinase signalling.

Overall, our proximity analyses of the *Leishmania* kinetochore detected 20 of the 25 kinetochore proteins predicted based on studies in *T. brucei^12^.* We did not detect KKT15, KKT22, KKT23, KKT25 and whilst we could detect KKIP1, it was not sufficiently enriched over the control sample to be classed as proximal to KKT2, KKT3 or KKT19. KKT10 and KKT19 are identical in the catalytic domain and only differ at the N-terminus; peptide coverage of the N-terminus allowed us to define KKT19 as a proximal protein. The novel component we identified, KKT26, is conserved in trypanosomatids and is essential for parasite growth. KKT26 is present at the kinetochore in G1/S, indicating that it is a member of the inner kinetochore. Whilst at the sequence level the kinetoplastid kinetochore appears quite distinct^11^, our proximity map of the kinetochore revealed several expected proteins and complexes normally associated with chromosome segregation in other eukaryotes, such as the chromosome passenger complex, condensin and cohesin, suggesting there are at least some common molecular processes governing the separation of chromosomes in *Leishmania^18,23^*.

Regulation of the kinetoplastid kinetochore by protein kinases is a largely unknown process, the study of which may illuminate unique mechanisms that have evolved in this group of protists. We used XL-BioID and the kinetochore protein kinase KKT3 as a bait to follow the kinetochore as it assembles during the cell cycle, discovering a set of proteins already present early in the cell cycle which may be analogous to the CCAN network in other eukaryotes^34^. Some of these kinetochore components such as KKT4 and KKT20 are recruited in increased amounts as the cell cycle proceeds and are likely important for attachment to the mitotic spindle^28^. A key advance in the presented method is the ability to track both proximal proteins and phosphosites at a complex, which allowed us to determine phosphosites whose levels altered much more dynamically than the parent protein and are thus likely key regulators of kinetochore assembly. Cell cycle regulated changes in kinetochore proteins and phosphosites have been characterized in a comprehensive study of *T. brucei^35^*, but such work only provides whole cell levels of proteins and phosphosites.

Chemical inhibition of protein kinases is an often used approach yielding valuable information on the relationship between protein kinases and networks of phosphorylation sites *in vivo*. We used AB1 inhibition of KKT10/KKT19 (CLK1/CLK2)^13^ to show that XL-BioID can be used to generate a spatially focussed view of inhibitor action, detecting potential substrates of these protein kinases. Work in the bloodstream form of *T. brucei* showed that AB1 caused dispersal of kinetochore localised KKT1, KKT2, KKT4^14^. In the procyclic (insect) form, depletion of KKT10/KKT19 did not appear to have an effect on kinetochore localisation of KKT1, KKT4 or other tested kinetochore proteins^16^. Our results from promastigote *Leishmania* therefore appear more in line with those from procyclic *T. brucei*. We anticipate that this combination of proximity phosphoproteomics and chemical inhibition will be a useful new strategy to discover *in vivo* protein kinase driven signalling pathways.

## Methods

### Cell culture

*Leishmania mexicana* (MNYC/BZ/62/M379) promastigotes were cultured at 25°C in HOMEM medium (modified Eagle’s medium) supplemented with 10% (v/v) heat inactivated fetal calf serum (GIBCO) and 1% Penicillin/Streptomycin (Sigma-Aldrich). CRISPR-cas9 edited parasites were supplemented with the appropriate selection antibiotics: BirA*::BDF5, KKT2::BirA*, BirA*::KKT3 or KKT19::BirA*: 4μg ml^−1^ puromycin (Invivogen), BDF5::mT and KKT3::mT: 4μg ml^−1^ puromycin, 10μg ml^−1^ blasticidin, LoxP flanked KKT24, KKT26: 4μg ml^−1^ puromycin, 10μg ml^−1^ blasticidin, 15μg ml^−1^ G418, 32μg ml^−1^ hygromycin B, mNG::KKT24, mNG::KKT26 KKT2::mCh: 4μg ml^−1^ puromycin, 10μg ml^−1^ blasticidin.

### Generation of CRISPR-cas9 edited lines

Primers used to generate sgRNA expression cassettes and repair cassettes were designed using http://www.leishgedit.net ^15^ or manually for the LoxP lines^25^. Cassettes were generated by PCR, precipitated with glycogen, resuspended in 10μl water for each transfection and heat sterilised for 5mins at 95°C. 2μl of this was then used to transfect 1×10^6^ promastigotes using a 4D nucleofector (Lonza). Parasites used for transfections were: Cas9 T7^15^ and DiCre Cas9 T7^36^. 18hrs after transfection, selection antibiotics were added and lines were cloned by limiting dilution. The KKT2::mCh line was generated by C-terminally tagging endogenous KKT2 in Cas9 T7 according to procedure outlined above but with puromycin selection on a population of parasites. The mNG::KKT24 and mNG::KKT26 lines were generated by endogenously tagging genes at the N-terminus in Cas9 T7 with blasticidin selection on a population. Lines were verified by western blot and/or diagnostic PCR. For loxP lines, clones were also screened for excision efficiency by diagnostic PCR of genomic DNA.

### Immunofluorescence

KKT2::mCh, mNG::KKT24 and mNG::KKT26 were washed once in PBS before fixation with 1% para-formaldehyde (PFA) for 10mins at RT. PFA was quenched with 0.5M Tris pH8.5 for 5mins at RT. 1×10^6^ parasites were prepared for each line. Parasites were allowed to adhere to microscope slides (SuperFrost, Thermo scientific) for 15mins, permeabilised with 0.5% Triton X-100 for 10mins and blocked with 5% FBS in PBS for 15mins. Anti-mCherry antibody (mCherry Monoclonal Antibody 16D7, Alexa Fluor 647, Invitrogen) at 1:50 in 5% FBS in PBS was added for 1hr at RT. Parasites were washed 3× in PBS for 5mins each wash before mounting in ProLong Diamond (Thermo scientific). Images were acquired on an inverted Zeiss AxioObserver with a 100× lens. Image Z-stacks were blind deconvolved with the Microvolution plugin for Fiji using 100 iterations.

### XL-BioID

BirA*::BDF5, KKT2::BirA*, BirA*::KKT3, KKT19::BirA* parasites were grown to 4 × 10^6^ ml^−1^ at which point biotinylation was initiated in 3 replicates of each line, by adding biotin to 150 μM for 18hrs. BDF5::mT and KKT3::mT parasites were grown to 2 × 10^6^ ml^−1^ then synchronised by adding hydroxyurea to 0.4 mg ml^−1^ for 18hrs^21^. Synchronised parasites were washed twice in culture medium and resuspended at 4 × 10^6^ ml^−1^ to proceed with the cell cycle, 5 replicates of each line and condition were used. For inhibition experiments, AB1 was added to a concentration of 60nM. Biotinylation time points were taken at 0, 4 and 8hrs after hydroxyurea wash off. At each time point, biotin was added to 0.5 mM for 30 mins biotinylation, 4 × 10^6^ parasites were taken for flow cytometry analysis.

After *in vivo* biotinylation, parasites were washed twice in PBS and resuspended to a density of 4 × 10^7^ ml^−1^ in pre-warmed PBS. DSP crosslinker was added to 1mM and *in vivo* cross-linking proceeded for 10mins at 25°C. Cross-linking was quenched for 5mins by addition of Tris-HCl pH7.5 to a concentration of 20mM. Parasites were harvested by centrifugation and pellets stored at −80°C until lysis. A pellet of 4 × 10^8^ parasites was used for each affinity purification which was lysed in 500 μl ice cold RIPA buffer containing 0.1 mM PMSF, 1 μg ml^−1^ pepstatin A, 1μM E64, 0.4mM 1-10 phenanthroline. In addition, every 10 ml of RIPA (0.1% sodium dodecyl sulfate, 0.5% sodium deoxycholate, 1% IgePal-CA-630, 0.1mM EDTA, 125mM NaCl, 50mM Tris pH7.5) was supplemented with 200μl proteoloc protease inhibitor cocktail containing w/v 2.16% 4-(2-aminoethyl)benzenesulfonyl fluoride hydrochloride, 0.047% aprotinin, 0.156% bestatin, 0.049% E-64, 0.084% Leupeptin, 0.093% Pepstatin A (Abcam), 3 tablets complete protease inhibitor EDTA free (Roche) and 1 tablet PhosSTOP (Roche). Lysates were sonicated with a microtip sonicator on ice for 3 rounds of 10 seconds each at an amplitude of 30. 1 μl of Benzonase (250 Units, Abcam) was added to each lysate, digestion of nucleic acids proceeded for 10 mins at RT followed by 50 mins on ice. Lysates were clarified by centrifugation at 10,000g for 10 minutes at 4°C. For enrichment of biotinylated material, 100μl of magnetic streptavidin bead suspension (1mg of beads, Resyn Bioscience) was used for each affinity purification from 4 × 10^8^ parasites. Biotinylated material was affinity purified by end-over-end rotation at 4°C overnight. Beads were washed in 500 μl of the following for 5 mins each: RIPA for 4 washes, 4 M urea in 50mM triethyl ammonium bicarbonate (TEAB) pH8.5, 6 M urea in 50 mM TEAB pH8.5, 1 M KCl, 50mM TEAB pH8.5. Beads from each affinity purification were then resuspended in 200 μl 50 mM TEAB pH8.5 containing 0.01% ProteaseMAX (Promega), 10 mM TCEP, 10 mM Iodoacetamide, 1 mM CaCl_2_ and 500 ng Trypsin Lys-C (Promega). On bead digest was carried out overnight at 37°C while shaking at 200 rpm. Supernatant from digests was retained and beads washed for 5 mins in 50 μl water which was then added to the supernatant. Digests were acidified with trifluoroacetic acid (TFA) to a final concentration of 0.5% before centrifugation for 10 mins at 17,000g. Supernatant was desalted using in house prepared C18 desalting tips, elution volume was 60 μl. Desalted peptides were either dried for MS analysis (BirA experiments) or used for enrichment of proximal phosphopeptides (miniTurboID experiments).

### Proximal phosphopeptide enrichment

Desalted peptides were pipette mixed to ensure a homogenous sample and 40% was removed and dried down for the ‘Total’ proximal protein sample. To the remaining 36 μl was added 51.2 μl acetonitrile, 10 μl 1 M glycolic acid and 5 μl TFA. Magnetic Ti-IMAC-HP beads (ReSyn Biosciences) were washed three times for 1 min in a 2× volume of loading buffer (0.1 M glycolic acid, 80% acetonitrile-ACN, 5% TFA). 10 μl of bead suspension (200 μg beads) were used to enrich phosphopeptides from each affinity purification. Peptides were added to beads and incubated with shaking at 800 rpm for 40 minutes at RT. Beads were washed for 2 mins at 800 rpm in 100 μl loading buffer, then 100 μl 80% ACN 1% TFA and finally 100 μl 10% ACN 0.2% TFA. Phosphopeptides were eluted in 2×40 μl 1% NH_4_OH for 10 minutes shaking at 800 rpm. 2 μl TFA was added and peptides dried down for MS analysis.

### Mass spectrometry data acquisition

For total proteome XL-BioID analysis peptides were loaded onto an UltiMate 3000 RSLCnano HPLC system (Thermo) equipped with a PepMap 100 Å C_18_, 5 μm trap column (300 μm × 5 mm, Thermo) and a PepMap, 2 μm, 100 Å, C_18_ EasyNano nanocapillary column (75 mm × 500 mm, Thermo). Separation used gradient elution of two solvents: solvent A, aqueous 1% (v:v) formic acid; solvent B, aqueous 80% (v:v) acetonitrile containing 1% (v:v) formic acid. The linear multi-step gradient profile was: 3-10% B over 8 mins, 10-35% B over 115 mins, 35-99% B over 30 mins and then proceeded to wash with 99% solvent B for 4 min. For analysis of samples prepared to measure dynamics of proximal proteins and phosphosites during kinetochore assembly, peptides were loaded onto an mClass nanoflow UPLC system (Waters) equipped with a nanoEaze M/Z Symmetry 100 Å C 18, 5 μm trap column (180 μm × 20 mm, Waters) and a PepMap, 2 μm, 100 Å, C 18 EasyNano nanocapillary column (75 mm × 500 mm, Thermo). Separation used gradient elution of two solvents: solvent A, aqueous 0.1% (v:v) formic acid; solvent B, acetonitrile containing 0.1% (v:v) formic acid. The linear multi-step gradient profile for ‘Total’ protein was: 3-10% B over 8 mins, 10-35% B over 115 mins, 35-99% B over 30 mins and then proceeded to wash with 99% solvent B for 4 min. For phosphopeptide enriched samples the following gradient profile was used: 3-10% B over 7 mins, 10-35% B over 30 mins, 35-99% B over 5 mins and then proceeded to wash with 99% solvent B for 4 min. In all cases the trap wash solvent was aqueous 0.05% (v:v) trifluoroacetic acid and the trapping flow rate was 15 μL/min. The trap was washed for 5 min before switching flow to the capillary column. The flow rate for the capillary column was 300 nL/min and the column temperature was 40°C. The column was returned to initial conditions and re-equilibrated for 15 min before subsequent injections.

The nanoLC system was interfaced with an Orbitrap Fusion hybrid mass spectrometer (Thermo) with an EasyNano ionisation source (Thermo). Positive ESI-MS and MS^2^ spectra were acquired using Xcalibur software (version 4.0, Thermo). Instrument source settings were: ion spray voltage, 1,900 V; sweep gas, 0 Arb; ion transfer tube temperature; 275°C. MS^1^ spectra were acquired in the Orbitrap with: 120,000 resolution, scan range: *m/z* 375-1,500; AGC target, 4e^5^; max fill time, 100 ms. Data dependent acquisition was performed in top speed mode using a fixed 1 s cycle, selecting the most intense precursors with charge states 2-5. Easy-IC was used for internal calibration. Dynamic exclusion was performed for 50 s post precursor selection and a minimum threshold for fragmentation was set at 5e^3^. MS^2^ spectra were acquired in the linear ion trap with: scan rate, turbo; quadrupole isolation, 1.6 *m/z*; activation type, HCD; activation energy: 32%; AGC target, 5e^3^; first mass, 110 *m/z*; max fill time, 100 ms. Acquisitions were arranged by Xcalibur to inject ions for all available parallelizable time.

### Mass spectrometry data analysis

Peak lists in .raw format were imported into Progenesis QI (Version 2.2., Waters) for peak picking and chromatographic alignment. A concatenated product ion peak list was exported in .mgf format for database searching against the *Leishmania mexicana* subset of the TriTrypDB (8,250 sequences; 5,180,224 residues) database, appended with common proteomic contaminants. Mascot Daemon (version 2.6.1, Matrix Science) was used to submit searches to a locally-running copy of the Mascot program (Matrix Science Ltd., version 2.7.0.1). Search criteria specified: Enzyme, trypsin; Max missed cleavages, 2; Fixed modifications, Carbamidomethyl (C); Variable modifications, Oxidation (M), Phospho (STY), Acetyl (Protein N-term, K), Biotin (Protein N-term, K); Peptide tolerance, 3 ppm (# 13C = 1); MS/MS tolerance, 0.5 Da; Instrument, ESI-TRAP. Peptide identifications were passed through the percolator algorithm to achieve a 1% false discovery rate as assessed empirically by reverse database search, and individual matches filtered to require minimum expect scores of 0.05. The Mascot .XML results file was imported into Progenesis QI, and peptide identifications associated with precursor peak areas were mapped between acquisitions. Relative protein abundances were calculated using precursor ion areas from non-conflicting unique peptides. For total proteome data, only non-modified peptides were used for protein-level quantification. Statistical testing was performed in Progenesis QI, with the null hypothesis being peptides are of equal abundance among all samples. ANOVA-derived p-values from QI were converted to multiple test-corrected q-values within QI. For phosphopeptide identifications, Mascot-derived site localisation probabilities were used. To associate quantification data with site localisation probabilities, Mascot search results including raw spectral details and QI quantification data were exported separately in .csv format. Empty rows in the Mascot .csv were removed in R (3.6.1) before joining the data from the two .csv files using KNIME Analytics Platform (4.3.1 KNIME AG). The combined .csv was stripped of non-quantified peptides before calculating Hochberg and Benjamini FDR q-values from the original QI ANOVA p-values.

For BirA* experiments, normalised protein label free peak areas were analysed with SAINTq^37^ using the following parameters: normalize_control=FALSE, compress_n_ctrl=100, compress_n_rep=100. False discovery rate for identified proximals was 1%.

For miniTurboID total proximal data, peptide ion quantification data was exported from Progenesis LFQ and missing value imputation was performed for each sample group, drawing values from a left shifted normal log2 intensity distribution to model low abundance proteins (mean=12.1, sd=1.5). For each protein, a mean intensity profile across all samples was calculated from peptide ion intensities. For each peptide ion, a pearson correlation coefficient was calculated between the mean protein intensity profile and the individual peptide ion intensity profile. Peptide ions with a correlation coefficient >0.4 were then summed to calculate the label free intensity of the parent protein. Protein intensities were log2 transformed and proximal proteins were determined with the limma package^38^ using options trend=TRUE and robust=TRUE for the eBayes function. False discovery rate for identified proximals was 1%.

For miniTurboID phosphosite proximal data, label free intensities were exported from Progenesis LFQ and missing values imputed by drawing values from a left shifted normal log2 intensity distribution to model low abundance phosphopeptides (mean=4, sd=1.2). Phosphosites were aggregated by summing intensities which were then log2 transformed. Proximal phosphosites were determined with the limma package^38^ using options trend=TRUE and robust=TRUE for the eBayes function. False discovery rate for proximal phosphosites was 5%.

## Data availability

Mass spectrometry data sets and proteomic identifications are available to download from MassIVE (MSV000087750), [<u>doi:10.25345/C5G543</u>] and ProteomeXchange (PXD027080).

## Acknowledgements

We thank Tony Larson for data analysis expertise and Rachel Neish for generating the DiCre Cas9 T7 line. This work was funded by the Wellcome Trust (200807/Z/16/Z). The research has also, in part, received funding from the Research Council United Kingdom Grand Challenges Research Funder under grant agreement ‘A Global Network for Neglected Tropical Diseases’ grant number MR/P027989/1. The York Centre of Excellence in Mass Spectrometry was created thanks to a major capital investment through Science City York, supported by Yorkshire Forward with funds from the Northern Way Initiative, and subsequent support from EPSRC (EP/K039660/1; EP/M028127/1).

## Contributions

VG and JCM conceived the project. JCM supervised the project. VG and JCM designed the experiments. VG and NGJ performed the experiments. VG and AD analysed experimental data. VG wrote the manuscript and JCM, NGJ and AD revised it.

## Competing interests

The authors declare no competing interests

## Supplementary Tables

Supplementary Table 1 - Comparison of BioID and XL-BioID proteomics with label free quantification.

Supplementary Table 2 - Proximal proteins associated with KKT2, KKT3 and KKT19 identified by Xl-BioID

Supplementary Table 3 - All proteins associated with KKT2, KKT3 and KKT19 identified by Xl-BioID

Supplementary Table 4 - All proteins associated with KKT3, including after AB1 treatment, identified by XL-BioID using miniTurbo.

Supplementary Table 5 - All phosphopeptides associated with KKT3, including after AB1 treatment, identified by XL-BioID using miniTurbo.

## Notes

https://massive.ucsd.edu/ProteoSAFe/dataset.jsp?accession=MSV000087750

http://proteomecentral.proteomexchange.org/cgi/GetDataset?ID=PXD027080

### Competing Interest Statement

The authors have declared no competing interest.

